# Hoogsteen base pairs increase the susceptibility of double-stranded DNA to cytotoxic damage

**DOI:** 10.1101/2020.05.24.113951

**Authors:** Yu Xu, Akanksha Manghrani, Bei Liu, Honglue Shi, Uyen Pham, Amy Liu, Hashim M. Al-Hashimi

**Affiliations:** Department of Chemistry, Duke University, Durham, NC, 27710, USA; Department of Biochemistry, Duke University School of Medicine, Durham, NC, 27710, USA

**Keywords:** DNA dynamics, sequencing, m^1^A, AlkB, echinomycin

## Abstract

As the Watson-Crick faces of nucleobases are protected in double-stranded DNA (dsDNA), it is commonly assumed that deleterious alkylation damage to the Watson-Crick faces of nucleobases predominantly occurs when DNA becomes single-stranded during replication and transcription. However, damage to the Watson-Crick faces of nucleobases has been reported in dsDNA *in vitro* through mechanisms that are not understood. In addition, the extent of protection from methylation damage conferred by dsDNA relative to single-stranded DNA (ssDNA) has not been quantified. Watson-Crick base-pairs in dsDNA exist in dynamic equilibrium with Hoogsteen base-pairs that expose the Watson-Crick faces of purine nucleobases to solvent. Whether this can influence the damage susceptibility of dsDNA remains unknown. Using dot-blot and primer extension assays, we measured the susceptibility of adenine-N1 to methylation by dimethyl sulfate (DMS) when in an A-T Watson-Crick versus Hoogsteen conformation. Relative to unpaired adenines in a bulge, Watson-Crick A-T base-pairs in dsDNA only conferred ~130-fold protection against adenine-N1 methylation and this protection was reduced to ~40-fold for A(*syn*)-T Hoogsteen base-pairs embedded in a DNA-drug complex. Our results indicate that Watson-Crick faces of nucleobases are accessible to alkylating agents in canonical dsDNA and that Hoogsteen base-pairs increase this accessibility. Given the higher abundance of dsDNA relative to ssDNA, these results suggest that dsDNA could be a substantial source of cytotoxic damage. The work establishes DMS probing as a method for characterizing A(*syn*)-T Hoogsteen base pairs *in vitro* and also lays the foundation for a sequencing approach to map A(*syn*)-T Hoogsteen and unpaired adenines genome-wide *in vivo*.

## Introduction

The Watson-Crick faces of nucleobases are tucked in the interior of the DNA double helix where they are largely inaccessible to solvent, shielded by Watson-Crick hydrogen bonding, and protected from endogenous and environmental agents that may cause various deleterious forms of alkylation damage(1–3). Yet alkylation damage to the Watson-Crick faces of nucleobases does occur in nature (4–9) and can result in base modifications (Fig. 1A) that prevent Watson-Crick pairing and block or interfere with DNA replication. A variety of damage repair enzymes have evolved to address these lesions (7,10), which if left unrepaired, can be highly cytotoxic and/or mutagenic (4,11).

**Figure 1.**
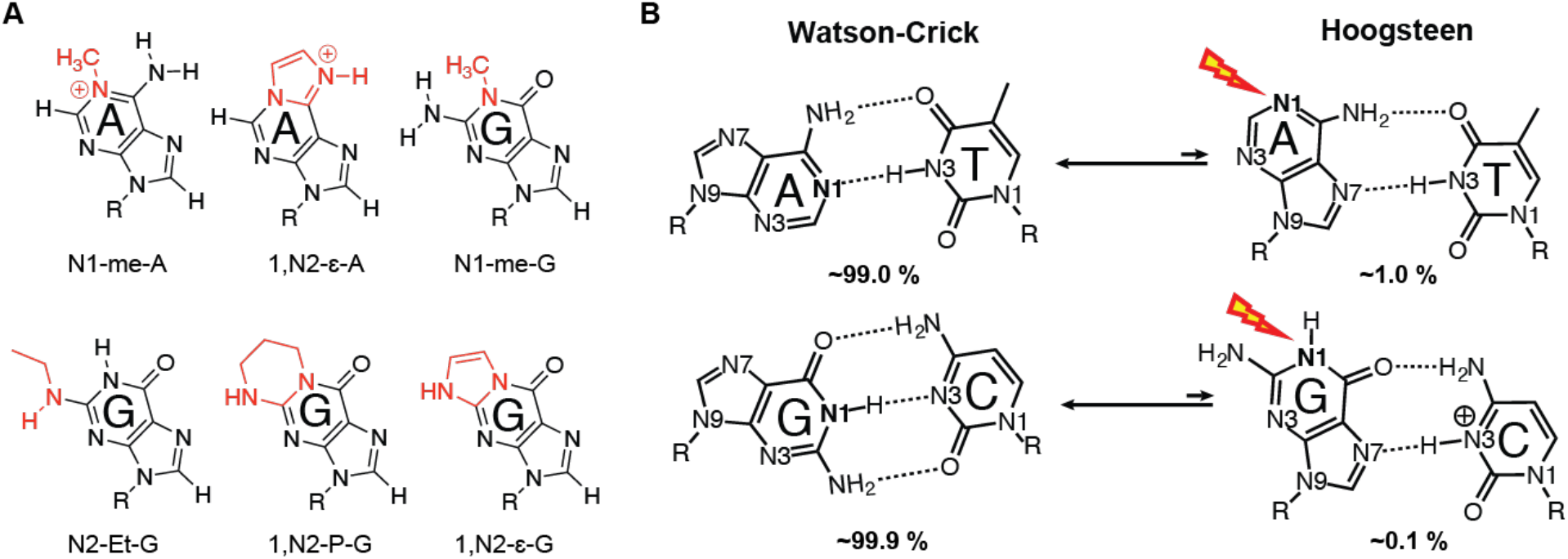
Proposed Hoogsteen-mediated alkylation damage to the Watson-Crick face of nucleotide bases. (A) DNA adducts targeting Watson-Crick faces of purine bases. (B) Exchange between Watson-Crick and Hoogsteen bps and proposed Hoogsteen-mediated alkylation damage to Watson-Crick face of purines.

It is generally accepted that alkylation damage to the Watson-Crick faces of nucleobases by endogenous and environmental agents as well as anti-cancer therapies (5,6,12) occurs primarily during replication and transcription, when the DNA is transiently single-stranded (4). *In vitro* studies have shown that Watson-Crick hydrogen bonding causes differences in the reactivity profiles of single-stranded (ssDNA) and doublestranded (dsDNA) DNA such that alkylation products at nitrogen functional groups involved in hydrogen bonding (e.g. adenine-N1 and cytosine-N3) are diminished in dsDNA relative to ssDNA, making adenine-N3 the most reactive site on the Watson-Crick face in dsDNA (9,13,14). Moreover, in prokaryotes, the activities of enzymes that repair alkylation damage have been linked to the process of DNA replication (4,15).

Based on prior studies, it is also clear that the Watson-Crick faces of nucleobases in dsDNA are indeed accessible to alkylation damage by reagents such as dimethyl sulfate (DMS)(13), methyl methanesulfonate (MMS)(14), ethylnitrosourea(9), and formaldehyde (16) *in vitro*. However, the mechanisms that underlie this phenomenon are poorly understood. Furthermore, the degree to which dsDNA protects Watson-Crick faces of nucleobases from alkylation damage relative to ssDNA has not been rigorously quantified. This is important given that the dsDNA is the dominant form of DNA *in vivo*, and that even if the reactivity were 1000-fold lower for dsDNA versus ssDNA, damage to dsDNA could still be a substantial source of alkylation damage as the abundance of the dsDNA exceeds that of ssDNA by an even greater amount.

In addition to becoming single-stranded, accessibility to the Watson-Crick face in duplex DNA has also been proposed to arise from alternative short-lived low-abundance conformational states that transiently expose nucleobases to solvent (9). It has long been established that in duplex DNA, Watson-Crick base pairs can spontaneously open and form conformations in which the otherwise buried and hydrogen bonded imino protons can exchange with solvent(17). Based on hydrogen exchange measurements *in vitro* in naked duplexes, the abundance of the base opened state is exceptionally low on the order of 10^−5^. A potentially more abundant conformational state that can increase the susceptibility of dsDNA to Watson-Crick face damage even further is the Hoogsteen conformation (18–21). A(*syn*)-T and G(*syn*)-C^+^ Hoogsteen bps form spontaneously in canonical dsDNA by flipping the purine bases 180° from an *anti* to *syn* conformation, leaving the Watson-Crick faces of the purine base exposed to solvent (Fig. 1B).

Hoogsteen-mediated alkylation damage to dsDNA could be substantial when considering that they form robustly across different DNA sequence and structural contexts with an abundance (0.1-5 %) that exceeds the base open conformation by at least two orders of magnitude (17). Based on *in vitro* measurements, there are millions of transient Hoogsteen bps in the human genome at any given time (20). Moreover, Hoogsteen bps can become the dominant conformation in DNA-protein and DNA-drug complexes (22–27). However, little is known regarding the vulnerability of dsDNA to damage when in the Hoogsteen conformation.

Here, we tested the hypothesis that adenine-N1 in the A(*syn*)-T Hoogsteen bp in dsDNA is more reactive to methylation by dimethyl sulfate (DMS) as compared to A(*anti*)-T Watson-Crick bps. In ssDNA, adenine-N1 is 10-fold more reactive than adenine-N3, and the major adenine DMS methylation product is m^1^A (28). In contrast, because adenine-N1 is protected in Watson-Crick bps in dsDNA, the major adenine DMS methylation product is m^3^A (14,28). Therefore, we hypothesized that A-T Hoogsteen bps in dsDNA would shift the reactivity from adenine-N3, which is exposed in both the Watson-Crick and Hoogsteen bps, to the more reactive adenine-N1, which is only exposed in the Hoogsteen conformation. In addition to providing insights into the potential role of Hoogsteen bps in DNA damage, this unique reactivity signature could also provide a means by which to discriminate Hoogsteen versus Watson-Crick bps *in vitro* and possibly *in vivo* using sequencing-based approaches. Using DMS as the methylating agent allowed us to investigate the alkylation susceptibility of Hoogsteen bps, and also to develop a methodology for detecting Hoogsteen bps given the well-established utility of DMS in mapping nucleic acid structure *in vitro* and *in vivo* (29–33).

Given that the N1-me-A (m^1^A) (Fig. 1A), the product of N1-methylation of adenine is a highly toxic lesion that blocks Watson-Crick pairing and DNA replication (4–6,12), a variety of repair enzymes have evolved to address this lesion. m^1^A is repaired by alphaketoglutarate-dependent dioxygenase (AlkB) (10,34,35) and its human analogs ABH2 and ABH3 (36,37) via oxidative demethylation. Our methodology for detecting m^1^A takes advantage of AlkB-mediated repair to enhance the specificity of m^1^A detection.

Our results reveal a mechanism for damaging Watson-Crick faces of DNA via A-T Hoogsteen bps without the need for melting dsDNA, and also establish the utility of DMS probing in characterizing A(*syn*)-T Hoogsteen base pairs in addition to unpaired adenines in dsDNA *in vitro*. This work also lays the foundation for a new sequencing approach to map Hoogsteen and unpaired nucleotides genome-wide *in vivo*.

## Results

### Assaying the specificity of the m^1^A antibody

We developed an antibody-based rescue-coupled dot-blot assay to specifically detect and quantify m^1^A following treatment of DNA oligonucleotides with DMS. Our assay integrates aspects of the m^1^A-MaP strategy used previously to map the m^1^A RNA methylome (38). It uses the anti-m^1^A monoclonal antibody (anti-m^1^A mAb, MBL International) (39) to specifically detect m^1^A. This antibody has previously been shown to specifically bind m^1^A and to discriminate against non-cognate nucleosides and nucleobases including m^6^A, m^1^G, m^2^G, m^7^G and unmethylated nucleosides, which bind with >1000-fold weaker affinity (39,40). The antibody has been shown to cross-react with the m^7^G-ppp-A extended cap structure in mRNA (41). However, this is of no consequence for our studies, which are focused on DNA oligonucleotides lacking this cap structure.

Prior studies have not tested the antibody specificity against m^3^A, which is a major adenine DMS methylation product in dsDNA. Due to its chemical instability, it is currently infeasible to obtain oligonucleotide containing m^3^A. We therefore used the m^3^A nucleobase in competition-based dot-blot assays to test the binding specificity of the antibody against the m^3^A. The same assay was also used to verify binding specificity against m^7^G, which is another major DMS product of dsDNA. In agreement with prior studies (39,40), the antibody discriminates against m^7^G to which it binds with > 625-fold weaker affinity relative to m^1^A (Fig. 2A). Although to a smaller degree, the antibody also discriminated against m^3^A to which it binds with >25-fold weaker affinity relative to m^1^A.

**Figure 2.**
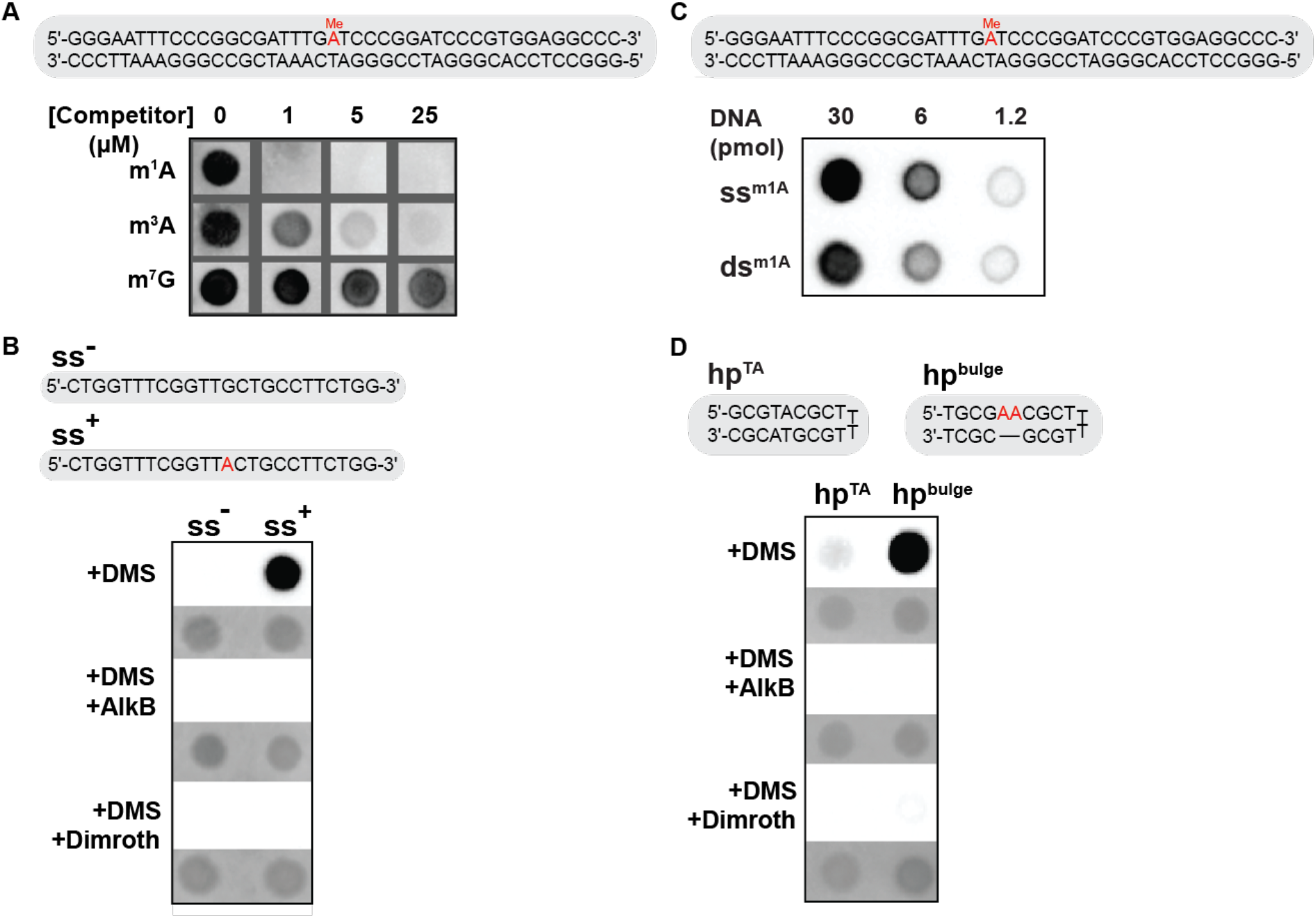
Sensitive detection of m^1^A using a dot-blot assay coupled to specific rescue. (A) Competition-based dot blot assay for assessing m^1^A antibody specificity. Shown are the dsDNA oligonucleotides used in the competition-based dot-blot assay along with the raw dot-blot data. m^1^A is highlighted in red. 50 pmol of the dsDNA was blotted on the membrane followed by incubation with the m^1^A antibody pre-mixed with the indicated amount of the competitor nucleobases. (B) Specific DMS induced m^1^A detection using rescue coupled dot-blot assay. Shown are the ssDNA oligonucleotides with (ss^+^) and without (ss^−^) a single adenine residue (highlighted in red) along with the raw dot blot data following 15 min DMS (75 mM) treatment pre- and post-AlkB repair and Dimroth reaction, along with their respective methylene blue loading controls (in grey). (C) DNA secondary structure independent detection of m^1^A. Shown are the DNA oligonucleotide containing m^1^A (highlighted in red) and the corresponding raw dot-blot data for ssDNA and dsDNA. (D) Specific DMS induced m^1^A detection at Watson-Crick AT bps and unpaired bulge adenines. Shown are the DNA oligonucleotides containing two adenines in bulge conformation (hp^bulge^) (highlighted in red) or in an A-T Watson-Crick (hp^TA^) along with the raw dot-blot data following 15 min DMS (75mM) treatment pre- and post-AlkB repair and Dimroth reaction. Also shown are their respective methylene blue loading controls (shaded grey).

To further test whether the anti-m^1^A mAb can discriminate m^1^A from other DMS products of ssDNA, including the major products m^7^G and m^3^C and the minor products m^3^G, O^6^-methyl-guanine, m^3^T and O^2^-methyl-thymine (28,42), we subjected (see Materials and Methods) ssDNA with (ss^+^) or without (ss^−^) a single adenine nucleotide to DMS treatment (Fig. 2B). As expected, ss^+^ showed a clear dot-blot signal following DMS treatment, while ss^−^ showed no detectable signal (Fig. 2B).

To further verify that the signal observed in ss^+^ is predominantly due to m^1^A and not m^3^A, we subjected the DMS-treated ss^+^ oligonucleotides to AlkB treatment. AlkB repairs m^1^A in both ssDNA and dsDNA but does not repair m^3^A (10,15,43). As expected, we did not observe any detectable signal from ss^+^ following AlkB treatment (Fig. 2B). As a final confirmation, we incubated the DMS-treated ss^+^ in a buffer and temperature condition that favors the Dimroth rearrangement (44,45), which specifically converts m^1^A, but not m^3^A, into m^6^A and observed no m^1^A signal in the dot-blot assay (Fig. 2B).

These results indicate that DMS methylates adenine-N1 in ssDNA, that the antibody enables detection of m^1^A with undetectable cross-reactivity to m^3^A or other DMS products under these conditions, and that AlkB treatment and the Dimroth rearrangement can be used to assess the specificity of m^1^A detection.

### The anti-m^1^A mAb antibody binds m^1^A in dsDNA

A prior study showed that an m^6^A antibody bound to m^6^A in double-stranded RNA with >10-fold weaker affinity as compared m^6^A in single-stranded RNA (46). We therefore compared the ability of the anti-m^1^A mAb antibody to recognize and bind m^1^A in dsDNA versus ssDNA. Unlike for m^6^A, we found the m^1^A antibody binds to m^1^A in dsDNA with only ~2-fold lower affinity relative to m^1^A in ssDNA (Fig. 2C). However, it should be noted that the relative affinity to ssDNA and dsDNA does vary with sequence context, with the affinity apparently becoming weaker for dsDNA when surrounding the m^1^A with stable G-C bps (data not shown).

### m^1^A detection following DMS treatment of bulge adenines

We benchmarked the AlkB rescue-coupled dot-blot assay by quantifying the m^1^A adduct following DMS treatment of two DNA duplexes containing two adenine residues (Fig. 2D) that are either exposed in a bulge (hp^bulge^) or protected by forming two canonical Watson-Crick A-T bps (hp^TA^). Hairpin constructs were used to increase duplex stability and minimize any reactivity from ssDNA due to melting or from having one strand in excess when using duplexes. A thymine rich apical loop was used to enhance the DNA crosslinking efficiency to the nylon membrane in the dot-blot assay (methods).

For these experiments, we sought to work under conditions of single-hit kinetics in which a given DNA molecule reacts no more than once with DMS (47–49). This was important for two reasons. First, this is necessary to ensure that the observed reactivity arises from the parent oligonucleotide and not from secondary methylation of singly methylated DNA. For example, production of m^7^G could destabilize a neighboring A-T bp, increasing its susceptibility to form m^1^A, and resulting in a spurious m^1^A signal. Second, duplexes with variable degrees of methylation may have different stabilities and this could also bias their ability to bind the antibody in the dot-blot assay. Having said that, any differences in reactivity due to either primary or secondary methylation seen for our highly controlled Watson-Crick versus Hoogsteen dsDNAs (see following sections) would have to originate from intrinsic differences in damage susceptibility between the Watson-Crick and Hoogsteen conformation.

Experiments were performed using 75 mM DMS for two reaction times (5 and 15 min) and the methylation stoichiometry was assessed using matrix-assisted laser desorption and ionization (MALDI) mass spectrometry (50). Based on the MALDI analysis of all DMS-treated samples, the major product for the 5 min reaction was singly methylated DNA whereas for the 15 min reaction, peaks corresponding to doubly methylated DNA were also observed (Fig. S1). We present results from both reaction conditions since the 15 min reaction time gave stronger signal-to-noise (S/N) and since the two data sets allows for more robust quantification of any differences in reactivity due to primary or secondary methylation between the Watson-Crick and Hoogsteen conformations. As will be detailed below, similar trends were observed for both reaction conditions.

In the dot-blot assay, a strong signal was observed for the DMS-treated hp^bulge^ indicating that the dinucleotide adenine bulge is accessible to DMS methylation (Fig. 2D). Moreover, no signal was detected post Dimroth reaction and following AlkB treatment, confirming that the signal primarily reflects the m^1^A product (Fig. 2D). Similar results were obtained for a different DNA sequence (Fig. S2). We also verified that the diminishment in m^1^A signal observed upon AlkB treatment results from the specific demethylation of m^1^A by performing the AlkB reaction in the absence of co-factors essential for catalytic activity and observed no detectable decrease in m^1^A signal (Fig. S2).

### DMS-treated dsDNA produces m^1^A likely through an alternative DNA conformation

Relative to the unpaired adenine bulge, the m^1^A signal was reduced substantially by as much as 130-fold for the Watson-Crick hp^TA^ duplex (Fig. 2D and Fig. S5). This residual signal is unlikely to arise from cross-reactivity with m^3^A or unmodified adenine, given that no signal was detected following AlkB treatment (Fig. 2D and Fig. S5), or from cross-reactivity with other bases and/or their DMS methylation adducts, given that no signal was detectable in the DMS-treated ss^−^ negative control (Fig. 2B).

Alternatively, the residual signal could correspond to m^1^A arising from methylating dsDNA, which has been reported previously (13). Here, our quantification of the signal allows us to consider alternative conformational states that could be susceptible to damage. The m^1^A signal was reduced by up to 130-fold for the hp^TA^ duplex relative to hp^bulge^. However, we would have expected ~100,000-fold reduction if m^1^A arose from methylating the base open state which has an abundance of ~10^−5^(17). Likewise, the signal is unlikely to arise from methylating the melted hairpin which is estimated to have an abundance of 10^−7^-10^−6^ (Table S1). The residual reactivity could arise from transient A-T Hoogsteen bps, which have an abundance of ~1%, assuming that the DMS reactivity of the adenine-N1 is similar in the Hoogsteen and bulge conformations. There could also be other as of yet poorly characterized conformational states in dsDNA that increase the susceptibility of adenine-N1 to methylation damage. Finally, while the residual signal was observed under single-hit kinetics conditions, we cannot entirely rule out that at least some of the signal reflects a minority of secondary methylated species which form in low-abundance under these reaction conditions and that fall outside the detection limit of MALDI.

### Echinomycin-DNA complexes as models for Watson-Crick and Hoogsteen base pairs

Assessing the reactivity of adenine-N1 in an A-T Hoogsteen bp requires the preparation of dsDNA samples containing the A-T Hoogsteen bp as the dominant conformation. To this end, we prepared complexes between dsDNA and echinomycin; an antibiotic with anti-tumor activity (51,52) that binds to CpG steps (<u>CG</u>) in dsDNA. Prior studies have shown that when two echinomycin molecules (or its close analog triostin A) bind to DNA sequences containing a TpA step sandwiched between two CpG binding-sites (d(<u>CG</u>**TA**<u>CG</u>)), the TpA step forms two tandem A-T Hoogsteen bps (25,53–57). Based on the crystal structure of the echinomycin-DNA complex (56) (see Fig. 7), adenine-N1 in the A-T Hoogsteen bps is solvent accessible, and not blocked by the bound echinomycin molecules, and therefore should be accessible to DMS methylation. While the complex only allows us to assess the reactivity of tandem A-T Hoogsteen bps, and not the Hoogsteen bps flanked by Watson-Crick bps that occur transiently in naked DNA duplexes, tandem Hoogsteen bps are common in DNA-protein and DNA-drug complexes (reviewed in (22)). As a negative control, we also examined a complex in which the TpA step is replaced by an ApT step, which retains two A-T Watson-Crick bps even following echinomycin binding (58).

We designed self-complementary hairpins (Fig. 3A) suitable for the dot-blot assay by elongating the core sequences of d(<u>CG</u>**TA**<u>CG</u>) and d(<u>CG</u>**AT**<u>CG</u>) previously shown to form Hoogsteen (25) and Watson-Crick (58) A-T bps respectively, upon echinomycin or triostin A binding. Again, a thymine rich apical loop was used to enhance the crosslinking efficiency to the nylon membrane (Fig. 3A).

**Figure 3.**
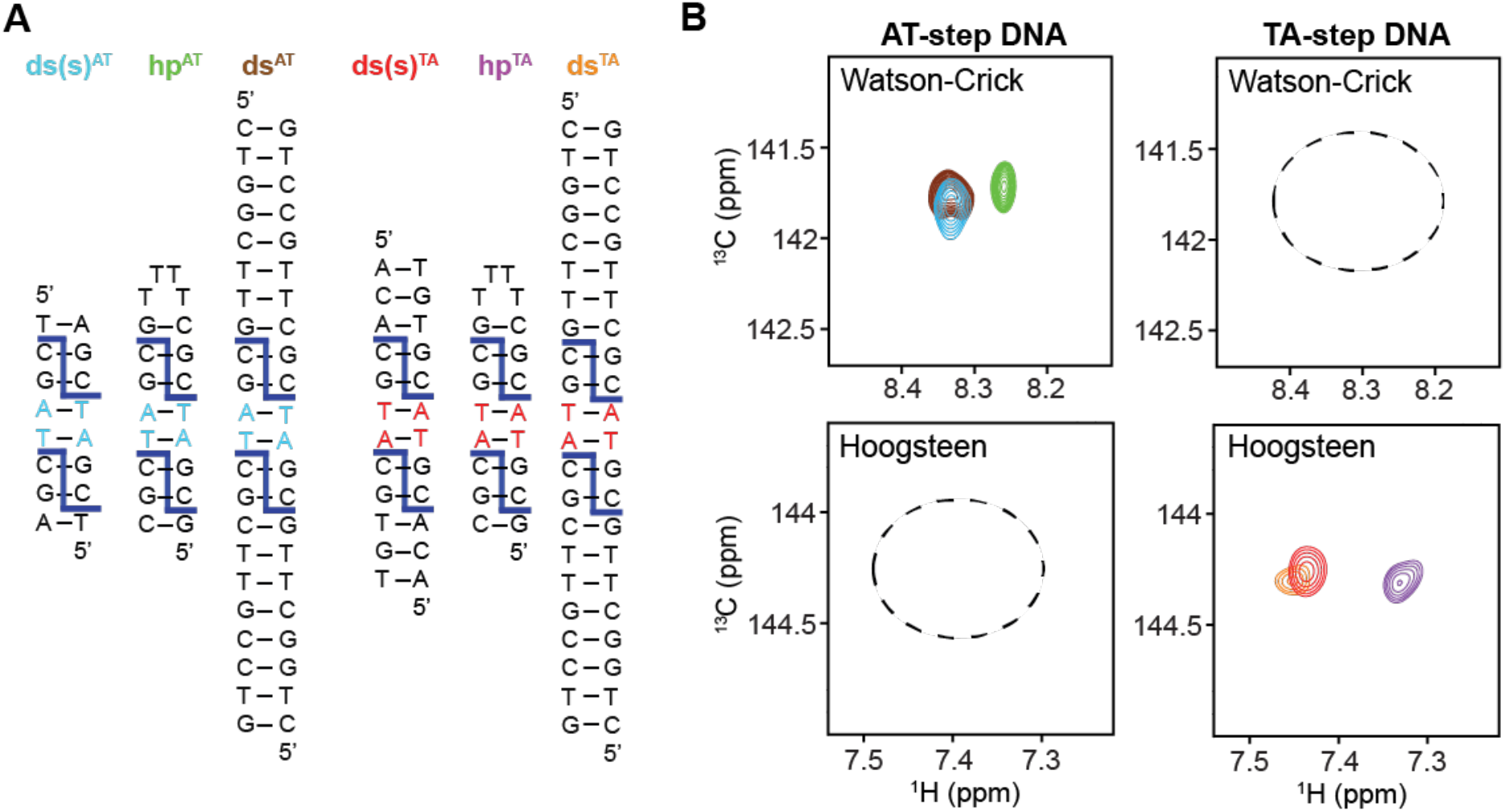
NMR experiments verifying formation of Hoogsteen and Watson-Crick bps upon drug binding. A) dsDNA oligonucleotides used in the NMR experiments. Shown are the reference dsDNA containing Watson-Crick (ds(s)^AT^) or Hoogsteen (ds(s)^TA^) bps, as well as hairpin (hp^AT^ and hp^TA^) and duplex (ds^AT^ and ds^TA^) dsDNA used in the dot-blot assays. Dark blue sticks represent the echinomycin molecule and its binding site at the CpG step. A-T bps adopting Hoogsteen or Watson-Crick conformation in the DNA-echinomycin complexes are highlighted in red and cyan, respectively. (B) Overlay comparing 2D [^13^C, ^1^H] HSQC NMR spectra of the aromatic region of DNA-echinomycin complexes for the reference dsDNA and the dsDNA used in the dot-blot assays.

Using NMR spectroscopy, we verified that the duplexes (Fig. 3A) do indeed bind to echinomycin as suggested by marked chemical shift perturbations in free DNA resonances upon intercalative binding of echinomycin (Fig. S3). To verify the A-T bp geometry in the TpA and ApT step complexes, we first prepared shortened versions of the duplexes (ds(s)^TA^ and ds(s)^AT^) previously shown by NMR (57,58) to form Hoogsteen and Watson-Crick bps, respectively upon echinomycin binding, in which only the TpA or ApT step was selectively labeled with uniformly ^13^C/^15^N nucleotides (Fig. 3A). We observed the expected chemical shift signatures unique to A-T Hoogsteen bps (59,60) in the TpA but not ApT step (Fig. 3B), including the downfield shifted adenine-C8 (144.3 ppm) accompanying the flip of the adenine into a *syn* conformation. Using these spectra as reference for the Hoogsteen and Watson-Crick conformations, we were able to verify formation of the Hoogsteen and Watson-Crick bps in the hp^TA^, hp^AT^, ds^TA^ and ds^AT^ sequences used in our dot-blot assay based on presence or absence of Hoogsteen chemical shift signatures (Fig. 3B).

### DMS-treated A-T Hoogsteen base pairs show enrichment in m^1^A relative to Watson-Crick base pairs

Next, we subjected the dsDNA oligonucleotides (Fig. 4A) to our dot-blot assay in the absence and presence of echinomycin. Following DMS treatment and quenching, the free DNA and DNA-echinomycin complexes were purified to remove residual β-ME, salt, and echinomycin. The methylated DNA was then divided into three aliquots; one aliquot was subjected to AlkB rescue, another to the Dimroth reaction, and a third aliquot was used for m^1^A quantification.

**Figure 4.**
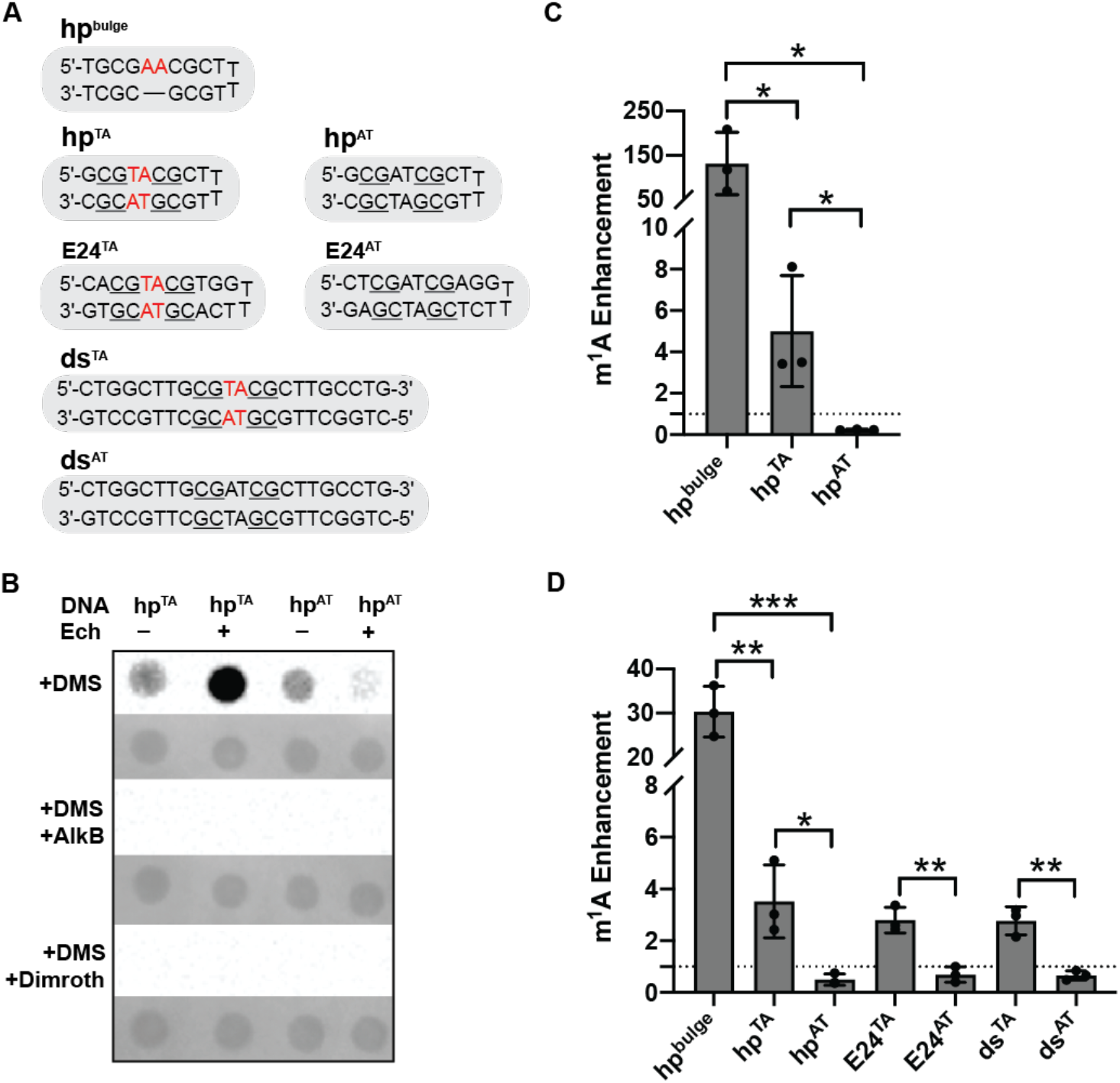
Dot-blot assay showing m^1^A enhancement in echinomycin induced A-T Hoogsteen bps. (A) The dsDNA oligonucleotides used in the dot-blot assay. The CG echinomycin binding sites are underlined. The unpaired adenines in the bulge and the A-T bps that convert into the Hoogsteen conformation upon echinomycin binding are shown in red. (B) A set of representative dot-blots of free and echinomycin bound DNA under 15 min DMS (75mM) treatment with and without AlkB repair and Dimroth reaction along with their respective methylene blue loading controls (shaded grey). (C) Quantification of dot-blot data for hp^bulge^, hp^TA^ and hp^AT^ under 5 min DMS (75 mM) treatment. (D) Quantification of dot-blot data for three sets of DNA sequences under 15 min DMS (75 mM) treatment. The bar plots in (C) and (D) show the enhancement of the m^1^A signal between the DNA-echinomycin complexes and their free DNA counterparts. For hp^bulge^, the m^1^A enhancement is calculated with respect to free hp^TA^ DNA. Shown are the average m^1^A enhancement from three independent DMS treatment replicates. The error bars represent the standard deviation. Statistical significance is calculated using the unpaired two-tailed parametric student’s t-test with 95% confidence interval. * denotes p < 0.05, ** denotes p < 0.01 and *** denotes p < 0.001.

Once again, the unbound hp^TA^ and hp^AT^ Watson-Crick DNA duplexes showed a small but detectable m^1^A signal (Fig. 4B). The DNA-echinomycin complexes showed markedly different m^1^A signal intensity than the unbound DNA (Fig. 4B). To assess the impact of echinomycin binding on reactivity, we computed echinomycin-induced m^1^A enhancement as the ratio between m^1^A signal of the dsDNA-echinomycin complex and its unbound dsDNA counterpart.

For the Watson-Crick hp^AT^ dsDNA, echinomycin binding reduced the m^1^A signal ~2-4-fold relative to the unbound hp^AT^ DNA (Fig. 4C-D). This could reflect quenching of transient Hoogsteen bps at these sites in the naked hp^AT^ duplex upon echinomycin binding. It was previously proposed (61) that the dipole interaction between the quinoxaline group of echinomycin and its adjacent A-T bp does not favor a Hoogsteen bp when an ApT step is sandwiched between echinomycin binding sites. Alternatively, the bound echinomycin may protect the bps neighboring the A-T and reduce secondary methylation of the A-T bps or even protect the A-T bps directly through non-specific binding, thereby reducing m^1^A relative to the free duplex. Any secondary methylation would have to be sufficiently low in abundance to remain undetected by MALDI.

In contrast, for the Hoogsteen hp^TA^ DNA, we observed ~4-fold echinomycin-induced m^1^A enhancement (Fig. 4C-D). Moreover, the enhancements were robustly observed across varying sequence contexts, in duplex and hairpin contexts, and for varying DMS incubation times (Fig. 4C-D and S4-S5). In all cases the signal was severely attenuated following AlkB treatment or the Dimroth reaction, confirming that the signal mainly corresponds to the m^1^A product. These results show that adenine-N1 is more susceptible to DMS methylation in the Hoogsteen versus Watson-Crick conformation.

Relative to Watson-Crick bps, the m^1^A enhancement in the echinomycin-induced A-T Hoogsteen bps under single-hit kinetic conditions was ~40-fold lower relative to the corresponding enhancement seen for unpaired bulge adenines in hp^bulge^ (Fig. 4C, compare hp^bulge^ with hp^TA^). This implies that adenine-N1 in an echinomycin-induced A-T Hoogsteen bp is 40-fold less reactive than unpaired adenine-N1. It is possible that the more constrained Hoogsteen conformation diminishes reactivity relative to unpaired adenines by sterically hindering the S_N_2 geometry required for DMS attack(13,62). If this were the case, the reactivity observed for naked Watson-Crick duplexes (see Fig. 2D, hp^TA^) could not be explained by the transient Hoogsteen bps. As described above, it is also possible that echinomycin protects the DNA from primary or secondary methylation in the drug complex and therefore results in reduction m^1^A relative to the free duplex, in which case the Hoogsteen reactivity in the complex underestimates its reactivity in naked DNA. If we assume that the reactivity of adenine-N1 in Hoogsteen is under-estimated by 4-fold given that echinomycin binding confers 2-4-fold protection to the Watson-Crick hp^AT^ DNA (Fig. 4C), then the Hoogsteen reactivity would only be 10-fold lower than the unpaired adenine. At this level of reactivity, it is possible that some of the m^1^A signal seen in the naked DNA duplexes reflects transient A-T Hoogsteen bps.

### AlkB-coupled primer extension reveals enrichment of m^1^A at Hoogsteen base pairs

To further confirm that the enhanced DMS reactivity seen in the echinomycin-hp^TA^ complex arises from methylation of adenine-N1 specifically at the A-T Hoogsteen bps, and to set the foundation for a new sequencing strategy to detect adenine nucleotides in unpaired and Hoogsteen conformations, we performed primer extension on DMS-treated duplexes that form either Watson-Crick or Hoogsteen bps upon echinomycin binding, pre- and post-AlkB repair.

For Watson-Crick bps in the unbound DNA duplexes and in the ds^AT-L^-echinomycin complex, the main DMS methylation product (m^3^A and m^7^G) are expected to induce AlkB-insensitive stops in the primer extension assay. m^3^A is a highly toxic lesion that blocks DNA replication (5), and although m^7^G generally does not affect DNA replication, it is prone to spontaneous depurination to form an abasic site that blocks DNA replication (63), and consequently it is typical to observe stops at unprotected guanines in primer extension assays (5,64). In contrast, for the tandem Hoogsteen A-T bps in the ds^TA-L^-echinomycin complex, the DMS methylation product m^1^A is expected to induce AlkB-sensitive stops.

Because of the lower sensitivity relative to the dot-blot assay, a higher DMS concentration (150 mM) and longer reaction time (15 min) needed to be used in the primer extension assay (33,65). While these conditions likely fall outside single-hit kinetics regime, the results from the dot-blot assay indicate that the relative trends in reactivity for the Hoogsteen and Watson-Crick bp are preserved between single- and multi-hit kinetics conditions.

As a positive control, we first verified that m^1^A efficiently terminates the Q5 High Fidelity DNA Polymerase (NEB Inc.) by subjecting two chemically synthetized dsDNA oligonucleotides with (ds^m1A^) or without (ds^unmod^) a single m^1^ A to the primer extension stop assay. As expected, while the major product for ds^unmod^ was full-length DNA, for ds^m1A^, the major product was DNA truncated immediately upstream of m^1^A (Fig. 5A). Furthermore, primer extension with AlkB-treated ds^m1A^ resulted in full-length DNA as the main product. Thus, m^1^A can efficiently terminate the primer extension of this DNA polymerase producing DNA products truncated one nucleotide before the m^1^A site.

**Figure 5.**
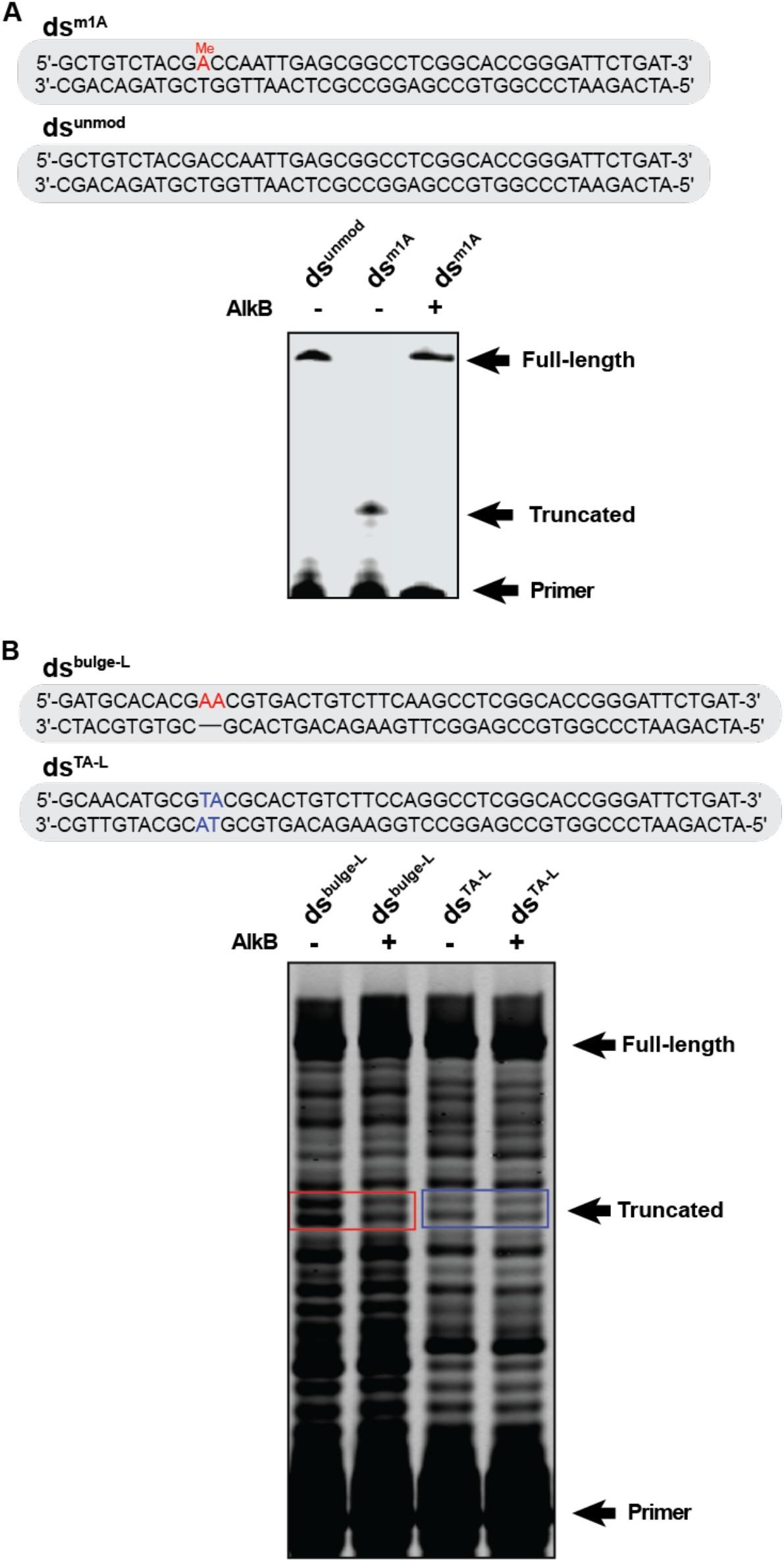
Primer extension stop assay on dsDNA containing m^1^A. (A) dsDNA oligonucleotides with (ds^m1A^) and without (ds^unmod^) m^1^A (highlighted in red) and the corresponding 10% urea polyacrylamide gel showing the primer extension products with and without AlkB repair. (B) DNA oligonucleotides containing either two unpaired adenines (ds^bulge-L^) (highlighted in red) or two A-T Watson-Crick bps (ds^TA-L^) (highlighted in blue) and the corresponding gel showing primer extension products of these DNA duplexes after 15 min DMS (150 mM) treatment with and without AlkB repair

To verify that the AlkB-coupled DMS primer extension assay can pinpoint unpaired adenines within the context of a dsDNA based on their unique adenine-N1 reactivity, we tested DNA duplexes similar to those used in the dot-blot assay containing either a dinucleotide adenine bulge (ds^bulge-L^, positive control) or two A-T Watson-Crick bps (ds^TA-L^, negative control) (Fig. 5B). For DMS-treated ds^TA-L^, the major primer extension product was the full-length DNA (Fig. 5B). Bands corresponding to products truncated at guanines and adenines could be observed when increasing exposure time during gel imaging. This is consistent with DMS-induced damage to Watson-Crick bps producing DNA replication blocking lesions m^7^G and m^3^A (64,66,67). As expected, these bands were insensitive to the AlkB treatment. DMS-treated ds^bulge-L^ showed a similar primer extension pattern as ds^TA-L^. However, two bands with 5-fold higher intensity as compared to ds^TA-L^ were observed corresponding to products truncated immediately before each of the two bulge adenines (Fig. 5B). These two bands were rescued following AlkB treatment, indicating that the polymerase stop can be attributed to m^1^A (Fig. 5B). These results establish the efficacy and robustness of the AlkB-coupled primer extension stop assay for detection of exposed adenine residues in duplex DNA at single nucleotide resolution.

Finally, we performed the primer extension assay on DMS-treated echinomycin-dsDNA complexes. The dsDNA sequences used in these studies were elongated (ds^TA-L^ and ds^AT-L^, see Fig. 6A) relative to those used in the dot-blot assay to allow for primer binding. For the unbound ds^TA-L^ and ds^AT-L^ duplexes, we observed the expected AlkB-insensitive products truncated immediately upstream of guanines and adenines (Fig. 6B). A similar pattern was also observed for the ds^AT-L^-echinomycin complex (Fig. 6B). In contrast, the band corresponding to the Hoogsteen site in the ds^TA-L^-echinomycin complex was highly sensitive to AlkB treatment (Fig. 6B, red box), indicating enrichment in m^1^A product at this site.

**Figure 6.**
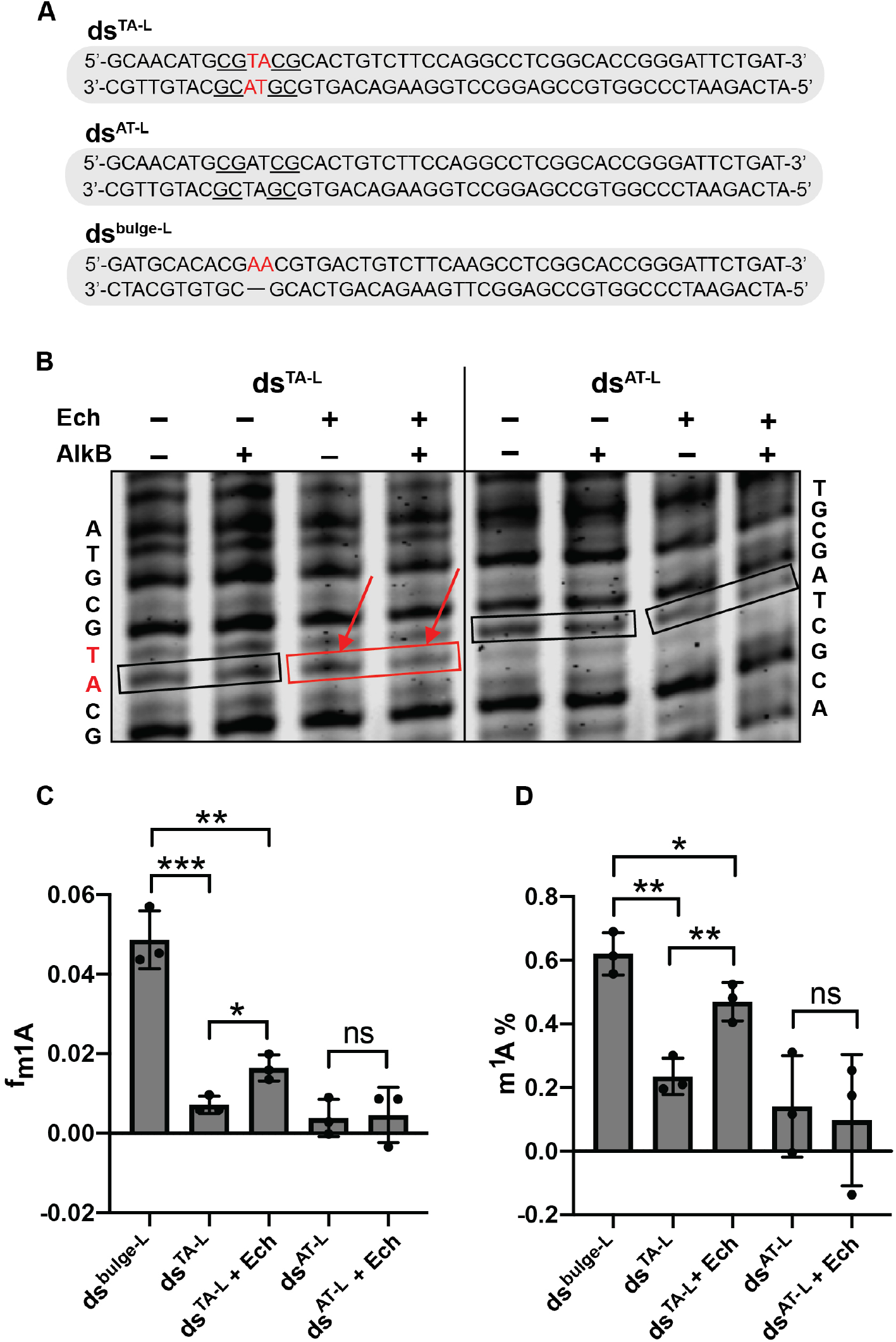
Primer extension stop assay on DMS-treated free and echinomycin bound DNA. (A) dsDNA oligonucleotides used in the primer extension assay. The CG echinomycin binding site is underlined. The unpaired adenines in the bulge and the A-T bps that convert into the Hoogsteen conformation upon echinomycin binding are shown in red. (B) A representative 10% urea polyacrylamide gel showing the products of primer extension for DMS-treated and AlkB-repaired free and echinomycin bound DNA. Bands corresponding to adenine of interest are highlighted with rectangles. Nucleotides at which the methylation causes the polymerase stop and thus giving the signal for a given band are indicated on the side of the gel. (C, D) Quantification of the m^1^A-induced polymerase stop frequency (f_m1A_) and the fraction of total stops attributable to m^1^A (m^1^A %) at the adenine of interest. The data represents average of three independent DMS treatment replicates and the uncertainty is reported as the standard deviation. Statistical significance is calculated using the unpaired two-tailed parametric student’s t-test with 95% confidence interval. ns denotes p > 0.05, * denotes p < 0.05, ** denotes p < 0.01 and *** denotes p < 0.001.

To quantify the m^1^A level in Hoogsteen versus Watson-Crick bps and compare band intensity across different samples, the band intensity of the truncation product at the adenine of interest was normalized relative to the band intensity of the full-length product in each sample. We define this normalized intensity as the polymerase stop frequency at each site due to m^1^A and m^3^A (f_m1A+m3A_). We then assigned the residual stops remaining following AlkB treatment to m^3^A (f_m3A_). The m^1^A-induced polymerase stop frequency (f_m1A_) is then given by f_m1A_ = f_m3A+m1A_ – f_m3A_.

As shown in Fig. 6C, the unbound free DNA (ds^TA-L^ and ds^AT-L^) showed some residual m^1^A signal, which is consistent with the results from the dot-blot assay (Fig 4B and S4A). The f_m1A_ value for the unpaired adenine in ds^bulge-L^ was 7-fold higher relative to the Watson-Crick A-T bp in unbound DNA, indicating that the unpaired adenine-N1 is 7-fold more reactive towards DMS as compared to adenine-N1 in a Watson-Crick bp under these reaction conditions. The 7-fold enhancement is in reasonable agreement with the ~30-fold (see Fig. 4D hp^bulge^) enhancement observed in the dot-blot assay for 15 min DMS reaction, especially when considering that a higher DMS concentration was used in the primer extension assay, which could disproportionately increase the apparent reactivity of the Watson-Crick duplex due to secondary methylation.

For the Hoogsteen A-T bp in the ds^TA-L^-echinomycin complex, the f_m1A_ was ~3-fold higher than the Watson-Crick A-T bp in its free DNA control (Fig. 6C), which is consistent with the dot-blot results for the 15 min DMS reaction (Fig. 4D). This provides additional support that the reactivity of adenine-N1 in the Hoogsteen versus Watson-Crick conformation does not vary considerably between single- and multi-hit kinetic conditions. In contrast, the ds^AT-L^-echinomycin complex did not show significant enhancement in m^1^A signal relative to the free DNA. Due to the low S/N, it was not possible to ascertain whether echinomycin binding confers a small degree of protection as observed in the dotblot assay (Fig. 6C). These results further confirm the higher reactivity of adenine-N1 in Hoogsteen versus Watson-Crick A-T conformations.

To compare the relative reactivity of adenine-N1 versus adenine-N3 in Watson-Crick versus Hoogsteen bps, we calculated the proportion of total stops attributable to m^1^A at the adenine of interest (m^1^A % = 1 - f_m3A_ / f_m3A+m1A_). As expected, for the naked DNA duplexes (ds^TA-L^ and ds^AT-L^), the majority (80%-90%) of the product was m^3^A whereas for the unpaired adenine bulge, the methylation reactivity shifted from adenine-N3 to adenine-N1, with ~60% of the product corresponding to m^1^A (Fig. 6D). This is consistent with the DMS methylation pattern observed in the ssDNA (28) and the canonical dsDNA (14), respectively. Relative to the Watson-Crick bps, the echinomycin– induced Hoogsteen in ds^TA-L^-echinomycin complex also shifted reactivity toward m^1^A, which accounted for 50% of the total adenine product, similar to that seen for unpaired adenines. In contrast, no statistically significant changes in reactivity were observed upon binding of echinomycin to ds^AT-L^. Thus, Hoogsteen bps shift the adenine DMS methylation patterns from m^3^A to m^1^A.

Taken together, these results show that Hoogsteen A-T bps increase the susceptibility of dsDNA to m^1^A damage, and the Hoogsteen sites can be identified as m^1^A sites in the AlkB-coupled primer extension stop assay at single nucleotide resolution.

## Discussion

It has been known for many decades that *in vitro*, the Watson-Crick faces of nucleobases in dsDNA are accessible to damage by alkylating reagents (9,13,14,16). Yet surprisingly, the extent of reactivity relative to ssDNA has not been quantitatively measured. Our results suggest that dsDNA only confers ~130-fold protection relative to unpaired nucleotides in a bulge. The reactivity could reflect formation of reactive transient Hoogsteen bps or some other unidentified conformational states. However, it could also arise as secondary methylation of dsDNA with the doubly methylated species falling below detection limits of MALDI. It could be that damage to solvent-exposed Hoogsteen faces of nucleobases to produce modifications such as m^7^G, could also increase the damage susceptibility of the Watson-Crick faces of nearby A-T bps, either by promoting Hoogsteen or other conformations. Further studies are needed to understand the origins of this dsDNA reactivity.

Damage to dsDNA either via alternative conformational states or as secondary products of damage can occur anytime during the cell life-cycle, and not only when the DNA is transiently single-stranded during replication and transcription. While studies have shown an association between AlkB repair activity and DNA replication in prokaryotes (15,68), no clear correlation between human ABH2 and ABH3 expression and cell proliferation has been established (4,36). Considering that the abundance of dsDNA greatly exceeds that of ssDNA, it remains plausible that in eukaryotes, adducts such as m^1^A accumulate in a DNA replication independent manner through damage to dsDNA.

Our results show that dsDNA is particularly vulnerable to damage when bps form the Hoogsteen conformation. For the echinomycin-induced tandem A-T Hoogsteen bps, the susceptibility of adenine-N1 to methylation was only 10-40 fold lower than unpaired adenine residues. Based on available crystal structures, similar solvent exposed A-T and G-C^+^ Hoogsteen bps can also be found in protein-DNA complexes such as the tumor suppressor protein p53 (27) and the general transcription factor TBP (26,69), which induce the A-T or G-C^+^ Hoogsteen bps, respectively, as the dominant conformation at specific sites (23,24,26,27,70–73) (Fig. 7). Some of these Hoogsteen bps have been verified under solution conditions (23,73). Structure modelling also reveals that a hypothetical A-T Hoogsteen bp forming in the context of the nucleosome core particle (74) would also expose adenine-N1 to solvent. Further studies are needed to investigate whether such Hoogsteen bps in protein-DNA complexes and possibly in other regions of the genome where DNA experiences torsional stress (75,76) are also vulnerable to Hoogsteen-mediated alkylation damage *in vitro* and *in vivo*.

**Figure 7.**
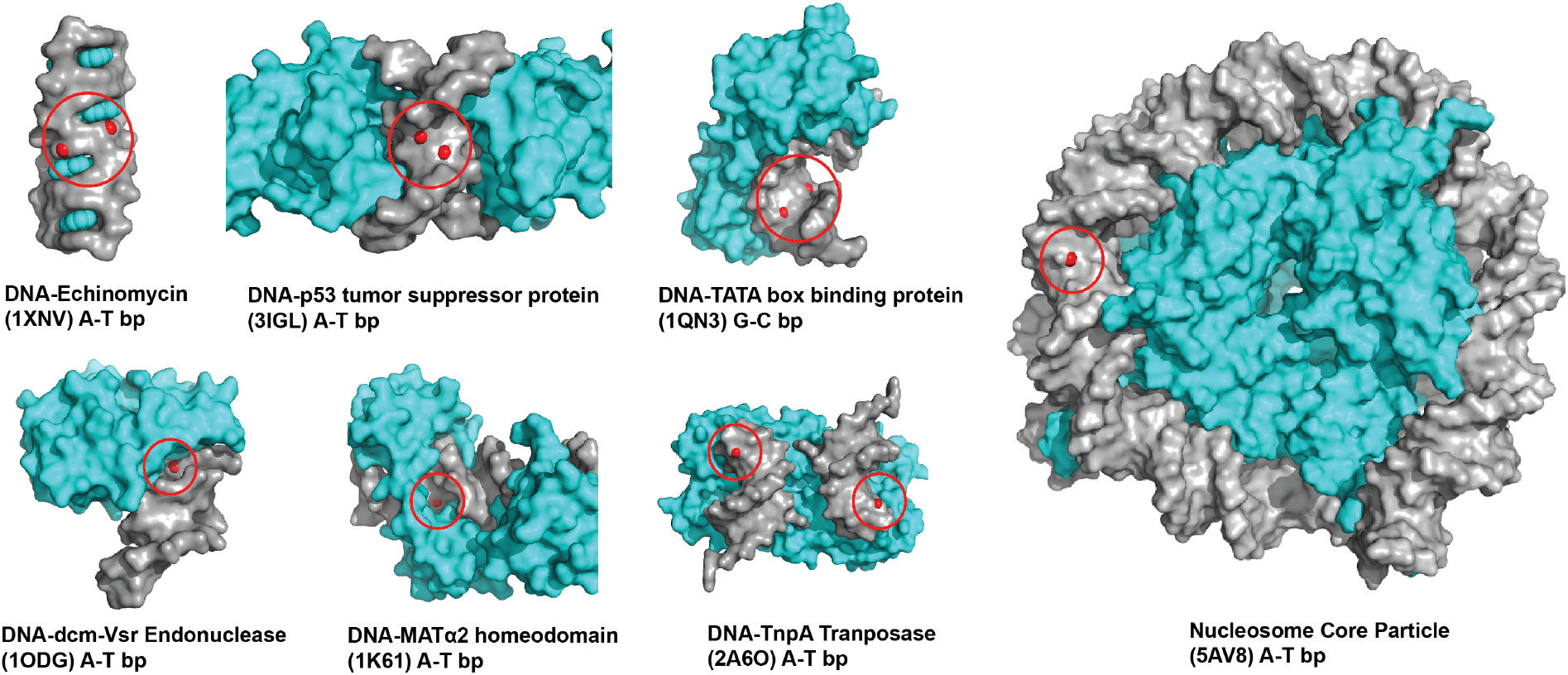
Solvent accessibility of purine-N1 in A-T and G-C Hoogsteen bps in crystal structures of DNA-protein and DNA-drug complexes. The surfaces of DNA and protein/drug are shown in grey and cyan, respectively. The adenine-N1 for A-T Hoogsteen or guanine-N1 for G-C^+^ Hoogsteen is shown in red and highlighted in a red circle. The structure of DNA-Integration host factor (1IHF) is not included above, as a subsequent solution state NMR study showed that the Hoogsteen bp was not formed under solution conditions (69). In the nucleosome core particle structure (PDB: 5AV8), residue DA-18 at chain I was manually flipped to a *syn* conformation using Pymol and then subjected to AMBER MD package(90) for energy minimization using AMBER ff99 force fields (91) as described previously(83).

The primer extension approach presented in this work provides a method for robustly identifying adenine nucleotides in an A-T Hoogsteen or unpaired conformation in protein-DNA complexes *in vitro* under solution conditions which can otherwise be difficult to visualize using other solution-state methods such as NMR (69). When combined with high-throughput sequencing, this approach could potentially be extended to map these structures genome-wide *in vivo* at single nucleotide resolution. In this regard, it is important to note that there have been many failed attempts to detect Hoogsteen bps in DNA-echinomycin complexes based on enhanced reactivity with a variety of reagents (66,77–79). However, none of these studies employed DNA sequences that were independently verified to contain Hoogsteen bps using techniques such as solution NMR. In addition, some of the chemical reagents used did not target sites that would be uniquely reactive in the Hoogsteen conformation (66,77,78). One study (80) showed that binding of echinomycin to DNA induced hyperreactivity to diethyl pyrocarbonate (DEPC) both at the echinomycin binding sites, as well as at a distal region containing alternating A-T bps. It was proposed that the hyperreactivity could result from A-T Hoogsteen bps, which would expose adenine-N3 in the major groove, and potentially cause enhanced reactivity to DEPC. It remains plausible that the reactivity seen by Mendel *et al* (80) did arise in part from enhanced transient Hoogsteen bps especially in light of recent NMR studies showing enhanced Hoogsteen breathing in an A-T rich region in a DNA-echinomycin complex (57). The DMS approach presented here could be applied to confirm the existence of enhanced transient Hoogsteen bps in these sequence contexts as well.

The high reactivity of adenine-N1 in the A-T Hoogsteen bp also has important implications for a growing number of studies that employ DMS to probe the secondary structure of RNA *in vitro* and *in vivo* (32,65,81,82). While it is typically assumed that high DMS reactivity corresponds to unpaired nucleotides, in RNA, Hoogsteen G-A and G-G mismatches with *syn* purine bases are common (69,83), and could be an additional source of high DMS reactivity in addition to unpaired nucleotides.

Finally, while we have focused on A-T Hoogsteen bps, G-C^+^ Hoogsteen bps could similarly provide mechanisms for inflicting damage to the Watson-Crick face of the guanosine base to produce many known forms of damage (Fig. 1A) including m^1^G, which occurs *in vivo* albeit in lower abundance than m^1^A and is highly mutagenic (84). It is interesting to note that the G-C^+^ Hoogsteen bp also occurs with ~10-fold lower abundance relative to the A-T Hoogsteen bp (20,21). While we have focused on alkylation of purine bases, Hoogsteen bps could also play roles in other forms of damage. For example, Hoogsteen bps have already been proposed to grant formaldehyde access to neighboring G-C Watson-Crick bps resulting in the hydroxymethylation of the cytosine amino nitrogen in dsDNA (85). Beyond Hoogsteen bps in dsDNA, other non-B DNA motifs have also been linked to mutagenesis that could expose Watson-Crick faces of nucleotide bases (86). Further studies are needed to understand how sequence-specific DNA dynamics may contribute to damage.

In conclusion, our results indicate that Watson-Crick faces of purine nucleobases in dsDNA are accessible to methylation by DMS and that Hoogsteen conformations increase this reactivity further. The unique reactivity of adenine-N1 in Hoogsteen and unpaired conformations provides a method to detect these conformations *in vitro* and possibly *in vivo*.

## Experimental procedures

### Reagents

DMS, 2-mercaptoethanol (β-ME), echinomycin, m^3^A and m^7^G nucleobases were purchased from Sigma-Aldrich. m^1^A monoclonal antibody (mouse) was purchased from MBL. The m^1^A nucleobase was purchased from Acros Organics. Horseradish peroxidase (HRP)-conjugated secondary antibody (anti-mouse IgG) was purchased from ThermoFisher. The AlkB enzyme was a gift from Dr. Patrick O’Brien (University of Michigan). Unmodified DNA oligonucleotides were purchased from IDT with HPLC purification for primer extension assays and using standard desalting for other experiments. 5’-IR700-labeled DNA oligos were purchased from IDT with HPLC purification. DNA oligonucleotides containing m^1^A or m^6^A were purchased from Yale Keck Biotechnology.

### DNA sample preparation

Duplex or hairpin dsDNA samples were prepared by dissolving oligonucleotides (1 mM for duplex, 20-100 μM for hairpin) in annealing buffer (15 mM sodium phosphate, 25 mM NaCl, 0.1 mM EDTA, and pH = 6.8). For non-palindromic duplexes used in the primer extension assay, 1x template strand was mixed with 1.5x complementary strand in order to eliminate any residual ss template strand. Samples were then heated at 95 °C for 10 min, followed by slowly cooling to room temperature for duplexes and rapid cooling on ice for hairpins. Samples were then buffer exchanged at least three times using a centrifugal concentrator (EMD Millipore, 3kDa cutoff) into the annealing buffer. Sample purity was then assessed based on the absorption ratios (A_260_/A_230_ and A_260_/A_280_) measured on NanoDrop 2000C (Thermo Fisher Scientific). The concentration of oligonucleotides was quantified using a Qubit fluorometer (High Sensitivity 1xdsDNA kit). DNA-echinomycin complexes were prepared by mixing the DNA in annealing buffer with 10x and 3x echinomycin dissolved in methanol for long (ds^TA-L^, ds^AT-L^) and short (hp^TA^, hp^AT^, ds^AT^, ds^TA^, E24^TA^ and E24^TA^) oligonucleotides, respectively, maintaining the annealing buffer:MeOH ratio at 2:1 (v:v). The complex solutions were incubated at room temperature for 45 min, followed by slow solvent evaporation under an air stream (55). The dried samples were re-dissolved in water, ensuring that the final salt concentration is identical to the annealing buffer.

### UV thermal denaturation

UV thermal denaturation experiments were performed on a PerkinElmer Lambda 25 UV/VIS spectrometer with an RTP 6 Peltier Temperature Programmer and a PCB 1500 Water Peltier System. DNA oligonucleotide stocks prepared in annealing buffer were diluted in annealing buffer to generate 3 μM 400 μL solutions. The samples were denatured at 1 °C/min with the absorbance at 260 nm (A_260_) being recorded every 0.5 °C. Three independent readings were recorded for each sample. Melting temperature (T_m_) and enthalpy change (ΔH^0^) for the melting transition were obtained by fitting the absorbance data to Equation (1) using an in-house Mathematica script:

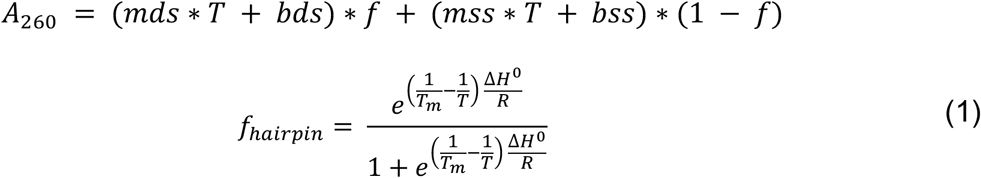

m_ds_ and b_ds_, mss and bss are coefficients representing the temperature dependence of the extinction coefficients of the hairpin and single strand, respectively. T is the temperature (units Kelvin), f is the fraction of the folded hairpin at a given temperature, T_m_ is the melting temperature (units Kelvin), ΔH^0^ is the enthalpy of the melting transition (units kcal/mol) and *R* is the universal gas constant (units kcal/K.mol). The entropy (ΔS^0^) and free energy (ΔG^0^) changes were computed from T_m_ and ΔH^0^ using Equation 2:

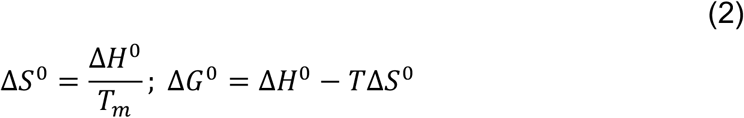

### DMS treatment

DNA samples (1000 and 100 pmol for the dot-blot and primer extension assays, respectively) in annealing buffer were treated with DMS (75 and 150 mM for the dot-blot and primer extension assays, respectively) at room temperature for 5 or 15 min. 2x volume of 66% ice-cold β-ME in DNA annealing buffer was added to quench the reaction. The DNA was immediately purified using a DNA cleanup column (the Monarch PCR & DNA cleanup kit from NEB) and eluted with water. The samples were stored at −20 °C until required for the next step in the protocol. Single-hit kinetics with respect to the DMS reaction was verified using mass spectrometry. Lack of over-methylation under the DMS treatment condition used was further confirmed by monitoring the bands corresponding to the full-length product in the primer extension stop assays, which were much stronger than those of the truncated products (Fig. S6).

### Matrix-assisted laser desorption/ionization–time of flight–mass spectrometry (MALDI–TOF–MS)

DMS-treated DNA oligonucleotides were desalted using a C18 Ziptip (Millipore). Briefly, the tip was primed with 50:50 H_2_O:MeCN with 0.1% TFA and rinsed with 0.1% TFA in H_2_O. 1μL sample was then loaded on the tip and desalted with 3×1 μL 100 mM ammonium acetate. The procedure was repeated two more times and the DNA oligo was finally eluted in 10:90 H_2_O:MeCN with 0.1% TFA. The MALDI matrix consisted of 30 mg/mL 3-hydroxypicolinic acid (3-HPA) and 10 mg/mL diammonium citrate (DAC) dissolved in 50:50 water:acetonitrile with 0.1% TFA. 1 μL matrix was first dried on the polished steel plate and then 1 μL of the desalted DNA oligo was dried over the top. Mass spectra were obtained using a Bruker Autoflex Speed LRF MALDI-TOF mass spectrometer equipped with a Nd:YAG laser (355 nm). Samples were analyzed in positive ion reflector mode using insulin/apomyoglobin as external calibration standards. Each spectrum was obtained using 500-1500 laser shots. The MALDI data of untreated hp^TA^ and hp^AT^ showed peaks corresponding to nucleobase losses that likely occur during MALDI sample preparation and/or desorption and ionization (87,88) as the number of resonances in the [^13^C,^1^H] 2D HSQC NMR spectrum (Fig. S3) is consistent with the base composition of the DNA duplex.

### AlkB repair

DMS-treated DNA samples (10 μL) were added to (90 μL) AlkB reaction buffer (25 mM HEPES, 100 mM NaCl, 2 mM sodium ascorbate, 1 mM 2-oxoglutarate, 40 μM (NH_4_)_2_Fe(SO_4_)_2_, 1 mM Tris(2-carboxyethyl)phosphone (TCEP), and 0.1 mg/mL BSA, pH 7.3). AlkB enzyme was then added to a final concentration of 2 μM. The reaction was allowed to proceed at 37 °C for 2h. Samples were then purified using the DNA cleanup column kit (the Monarch PCR & DNA cleanup kit from NEB). The specificity of AlkB repair reaction was confirmed using an inactive AlkB control reaction in which Fe^2+^, ascorbate and 2-oxoglutarate co-factors required for the demethylation reaction were not added (Fig. S2).

### Dimroth reaction

DMS-treated DNA samples (10 μL) were added to 1 mL 0.1 M Na_2_CO_3_/NaHCO_3_ buffer (pH 10.2) and incubated at 65 °C for 3h. Samples were then purified using the DNA cleanup column kit (the Monarch PCR & DNA cleanup kit from NEB).

### Dot-blot assay

A chemically synthesized oligonucleotide containing a single m^1^A was used to calibrate the m^1^A concentration range over which the m^1^A antibody signal intensity varies linearly with m^1^A concentration (Fig. S7). Based on this m^1^A standard curve, for the subsequent dot-blot assays, 100 pmoles of DNA samples were blotted on a positively charged nylon membrane (Amersham Hybond-N+, GE Healthcare Life Science). The DNA was then UV crosslinked to the membrane (254 nm, 15 min). The membrane was incubated in 0.03 % methylene blue (in 0.5 M acetate) for 30 min followed by a brief wash with PBST buffer (0.1% Tween 20 in 1x PBS, pH 7.4), and was then imaged to quantify the amount of DNA loaded on the membrane (loading controls). Methylene blue destaining was performed by washing with ethanol for 5 min. The membrane was blocked with 5% nonfat dry milk (in PBST buffer) at room temperature for 1h, and then incubated with anti-m^1^A mAb (1:5000 dilution in PBST buffer) at 4°C overnight followed by 3 x 10 min PBST washes. The membrane was then incubated with HRP-conjugated secondary antibody (1:2500 dilution in PBST buffer) at room temperature for 1h followed by 3 x 10 min PBST washes. The membrane was developed using enhanced chemiluminescent substrate (GE Healthcare) on Bio-Rad Chemi-Doc imager. The signal intensity was quantified using Image Lab software. For nucleobase-based competition assays, the anti-m^1^A mAb (1:5000 in PBST buffer) was pre-mixed with a series of concentrations of nucleobases (m^1^A, m^3^A or m^7^G) prior to incubation with the membrane containing m^1^A-incoporated dsDNA.

### Primer extension stop assay

Primer extension reactions were performed by mixing DNA oligonucleotide samples (0.1 μM x 2 μL in H_2_O) with the 5’-IR700 labeled primer (5’-ATCAGAATCCCGGTGCCGAGGC-3’) (1.5 μM x 2.5 μL in H_2_O) followed by addition of 4.5 μL NEBNext Ultra II Pol master mix (NEB). The reaction mixture was denatured at 98 °C for 1 min, then annealed and extended at 72 °C for 6 min, finally cooled to room temperature in a PCR machine (eppendorf AG). 8.5 μL of stop solution (0.05% orange G and 20 mM EDTA in formamide) was then added. Samples were denatured at 98 °C for 3 min and loaded immediately (7.5 μL) on a denaturing polyacrylamide gel (10% polyacrylamine, 8M Urea). The gel was run at 45°C and 45W for 50 min and visualized using the LI-COR odyssey Clx imaging system at 700 nm. The band intensities were quantified using Image Studio software.

### NMR experiments

1D and 2D NMR spectra were collected on a 700 MHz Bruker Avance III spectrometer equipped with a triple-resonance HCN cryogenic probe. Data were processed and analyzed with NMRpipe(89) and SPARKY (T.D. Goddard and D.G. Kneller, SPARKY 3, University of California, San Francisco), respectively.

## Acknowledgements

We thank Dr. Patrick O’Brien for kindly providing the AlkB enzyme. We thank the Duke Magnetic Resonance Spectroscopy Center for their technical support and resources. We thank Dr. Peter Silinski (Duke Chemistry Shared Instrument Facility) for performing the MALDI experiments (funded by North Carolina Biotechnology Center Grant #2017-IDG-1018).We also thank Atul K. Rangadurai and Megan Kelly for critical input.

## Funding information

This work was supported by a Mathers Foundation grant to H.M.A.

## Data Availability Statement

All data presented are available upon request from Hashim M. Al-Hashimi

## Conflict of interest

The authors declare that they have no conflict of interest with the contents of this article.

